# MicroPhenoDB Associates Metagenomic Data with Pathogenic Microbes, Microbial Core Genes, and Human Disease Phenotypes

**DOI:** 10.1101/2020.07.29.221010

**Authors:** Guocai Yao, Wenliang Zhang, Minglei Yang, Huan Yang, Jianbo Wang, Haiyue Zhang, Lai Wei, Zhi Xie, Weizhong Li

## Abstract

Microbes play important roles in human health and disease. The interaction between microbes and hosts is a reciprocal relationship, which remains largely under-explored. Current computational resources lack manually and consistently curated data to connect metagenomic data to pathogenic microbes, microbial core genes, and disease phenotypes. We developed the MicroPhenoDB database by manually curating and consistently integrating microbe-disease association data. MicroPhenoDB provides 5677 non-redundant associations between 1781 microbes and 542 human disease phenotypes across more than 22 human body sites. MicroPhenoDB also provides 696,934 relationships between 27,277 unique clade-specific core genes and 685 microbes. Disease phenotypes are classified and described using the Experimental Factor Ontology (EFO). A refined score model was developed to prioritize the associations based on evidential metrics. The sequence search option in MicroPhenoDB enables rapid identification of existing pathogenic microbes in samples without running the usual metagenomic data processing and assembly. MicroPhenoDB offers data browsing, searching and visualization through user-friendly web interfaces and web service application programming interfaces. MicroPhenoDB is the first database platform to detail the relationships between pathogenic microbes, core genes, and disease phenotypes. It will accelerate metagenomic data analysis and assist studies in decoding microbes related to human diseases. MicroPhenoDB is available through http://www.liwzlab.cn/microphenodb and http://lilab2.sysu.edu.cn/microphenodb.

## Introduction

The human body feeds a large number of microbes, mainly composed of bacteria, followed by archaea, fungi, viruses, and protozoa. Microbes, inhabiting various organs of the human body, mainly in gastrointestinal tract, as well as in respiratory tract, oral cavity, stomach, and skin, play important roles in human health and disease [1–3]. Microbial gene products have rich biochemical and metabolic activities in the host [4–6]. Microorganisms usually form a healthy symbiotic relationship with the host. However, when the microbial content becomes abnormal, or exogenous microbes infect the host, the balance of host microecology can be broken, which in turn can possibly cause various diseases [7, 8]. A tripartite network analysis in patients with irritable bowel syndrome demonstrated that the gut microbe *Clostridia* is significantly associated with brain functional connectivity and gastrointestinal sensorimotor function [9]. Strati *et al*. reported that Rett syndrome is substantially associated with a dysbiosis of both bacterial and fungal components of the gut microbiota [10]. The alteration of microbial communities on psoriatic skin is different from those on healthy skin and has a potential role in Th17 polarization to exacerbate cutaneous inflammation [11]. The ongoing pandemic of coronavirus disease 2019 (COVID-19) has affected more than 180 countries worldwide by April 2020. Lung injury has been reported in most patients with confirmed severe acute respiratory syndrome coronavirus 2 (SARS-CoV-2) infection [12].

The interaction between microbes and hosts is a reciprocal relationship and remains largely under-explored [13]. Accurate relationship information between microbes and diseases can greatly assist studies in human health [14]. With the wide application of next generation sequencing (NGS) technology, microbiological analysis methods and standards are being rapidly developed, such as metagenomic approaches [15]. As a result, a large amount of experimental data has been published [16], thus accurate database platforms are greatly needed to utilize these experimental data, determine the composition of pathogenic microbes in hosts, clarify microbial-disease relationships, and provide standardized high-quality annotation for clinical uses [17].

Due to the functional and clinical significance of microbes, several public databases have been established to collect microbe-disease association data, such as the Human Microbe-Disease Association Database (HMDAD) [18], Disbiome [19], the Virulence Factor Database (VFDB) [20], and the Comprehensive Antibiotic Resistance Database (CARD) [21]. HMDAD and Disbiome collate text-mining-based microbe–disease association data from peer-reviewed publications and describe the strength of the associations based on the credibility of the data sources. VFDB provides up-to-date knowledge of the virulence factors (VFs) of various bacterial pathogens; CARD contains high quality reference data on the molecular basis of antimicrobial resistance with an emphasis on genes, proteins, and mutations involved. Data in VFDB and CARD help to explain the relationship between pathogenic microbial genes and the health status of hosts. In addition, to assist physicians and healthcare providers to quickly and accurately diagnose infectious diseases in patients, a guideline for utilization of the microbiology laboratory for diagnosis of infectious diseases was developed and is being regularly updated by the Infectious Diseases Society of America (IDSA) and the American Society for Microbiology (ASM) [22]. The curation and analysis of microbe-disease association data are essential for expediting translational research and application. However, these computational resources lack manually and consistently curated data to connect metagenomic data to pathogenic microbes, microbial core genes, and disease phenotypes.

To bridge this gap, we developed the MicroPhenoDB database (http://www.liwzlab.cn/microphenodb) by manually curating and consistently integrating microbe-disease association data. We collected and curated the microbe-disease associations from the IDSA guideline [22], the National Cancer Institute (NCI) Thesaurus OBO Edition (NCIT) [23], and the HMDAD [18] and Disbiome [19] databases, and also connected microbial core genes derived from the MetaPhlAn2 dataset [24] to pathogenic microbes and human diseases. A refined score model was adopted to prioritize the microbe-disease associations based on evidential metrics [18, 25]. In addition, a sequence search web application was also implemented to allow users to query sequencing data to identify pathogenic microbes in metagenomic samples, as well as to retrieve the disease-related information of virulence factors and antibiotic resistances. MicroPhenoDB allows users to browse, search, access, and analyze data through the user-friendly web interfaces, visualizations and web service application programming interfaces (APIs).

## Data collection and processing

### Data collection and manual annotation

To ensure data quality, we not only integrated the association data with annotations from HMDAD and Disbiome, but also manually collated and curated microbe-disease association data from the IDSA guideline and NCIT (**Figure 1**). The IDSA guideline provides criteria for clinical identification of infectious microbes, while NCIT is a reference terminology that provides comprehensive information for infectious microbes. To enrich the annotation for disease-microbe associations, we manually traced the relevant literature in HMDAD and Disbiome; we also provided the microbes with annotation at the resolution of species level, such as taxonomies and official names. Association data between infectious microbes and diseases in IDSA were extracted. Relevant information about disease phenotypes and microbes in the microorganism notes from NCIT were extracted as well. The collected and integrated association data include information about microbe symbols, disease symbols, the increased or decreased impacts of the microbes, PubMed identifiers, and validation methods.

**Figure 1.**
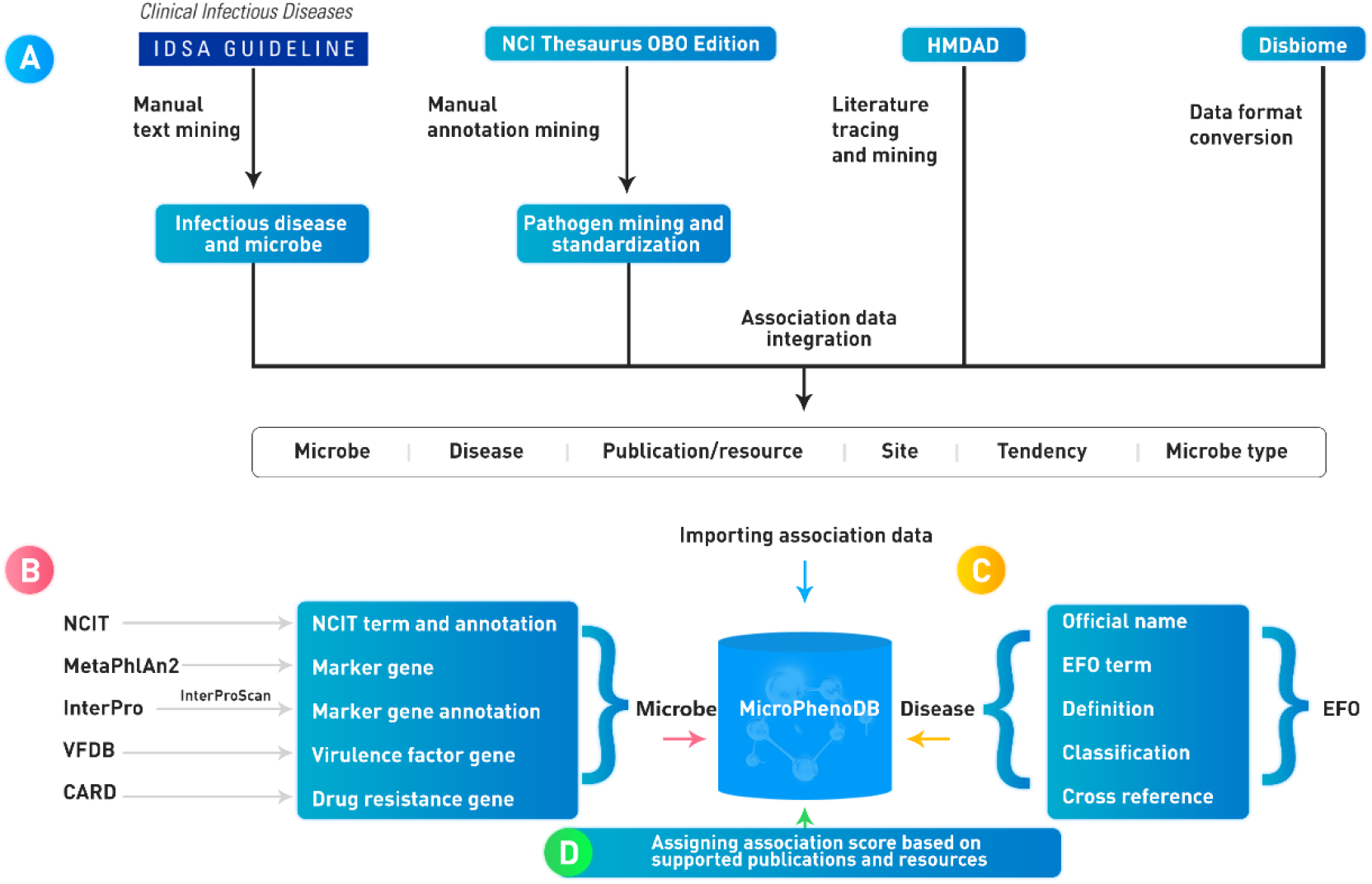
Workflow demonstrating the construction and curation of the MicroPhenoDB database

### Controlled vocabulary and ontology to describe microbes and diseases

In MicroPhenoDB, several resources of standard terminology and controlled vocabulary were adopted to consistently annotate microbes and diseases (Figure 1). Different tools and reference databases might give different taxonomies for microbes. To avoid this discrepancy, the official names of microbes were taken from NCIT [23] and the taxonomy identifiers were adopted from the National Center for Biotechnology Information (NCBI) [26] and UniProt [27]. The relationships between core genes and microbes were annotated using the MetaPhlAn2 tool [28], the microbial gene functions were annotated using the InterProScan tool [29], and the virulence factors and the drug resistance information of microbes were retrieved respectively from the databases of VFDB [20] and CARD [21]. The disease phenotypes were annotated with official names, experimental factor terms, definitions, classifications, and cross references using EFO [30]. EFO provides a systematic description of many experimental variables across European Bioinformatics Institute (EMBL-EBI) databases and the National Human Genome Research Institute (NHGRI) genome-wide association study (GWAS) catalog [31]; it also combines parts of several popular ontologies, such as Orphanet Rare Disease Ontology [32], Human Phenotype Ontology [33], and Monarch Disease Ontology [34]. The version or release of databases and tools used in the MicroPhenoDB construction are detailed in Supplementary Table 1.

### Association score model

One of the main problems in exploiting large collections of aggregated microbiome data is how to prioritize the associations. According to the previous studies by Ma *et al* [18] and Pinero *et al* [25], we refined the association score model to prioritize the microbe-disease associations using extra evidential metrics, including the number of sources that report the association, the type of curation of each source, and the number of supporting publications in the manual curation.

For every disease *i* and every microbe *j*, the raw score of their relationship *Raw_score_ij_* was defined as:

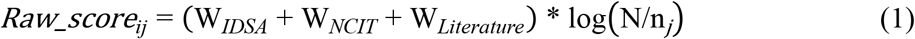

In Equation (1), *W_IDSA_* is the weight of the association source from the IDSA guideline, *W_NCIT_* is the weight of the association source from NCIT, and *W_Literature_* is the weight of the association source from literature publications. *N* is the number of all diseases in MicroPhenoDB, and *n_j_* is the number of diseases associated with microbe *j*. *Log*(*N/n_j_*) is computed to increase *Raw_score_ij_* for the microbes that are specifically associated with few diseases, or to decrease *Raw_score_ij_* for the microbes that are globally associated with several diverse diseases.

In Equations (2)-(4), MicroPhenoDB assigns different weights to different evidential sources according to their reliabilities (**Table 1**) [25]. If the association is curated from literature publications, *W_Literature_* is originally assigned as 0.25, otherwise assigned as 0. If the association is curated from NCIT [23], *W_NCIT_* is originally assigned as 0.5, which is double that of *W_Literature_*, otherwise assigned as 0. If the association is curated from IDSA [22], *W_IDSA_* is originally assigned as 1.0, which is double that of *W_NCIT_*, otherwise assigned as 0. The three weights also depend on the direction of the abundance change of a microbe in a disease and the number of supporting publications. *D_ij_* (*D_ij_* ∈ {1, −1}) represents the direction of the abundance change of microbe *j* in disease *i*. If the microbe *j* is increased in the case of disease *i*, *D_ij_* equals 1; if the microbe *j* is decreased in the case of disease *i*, *D_ij_* equals −1. *n_p_* is the number of publications in which an association between a disease and a microbe has been reported. From the distribution of numbers of evidences, we found *n_p_* was less than 16 and mostly ranged from 1 to 2 (Supplementary Figure 1).

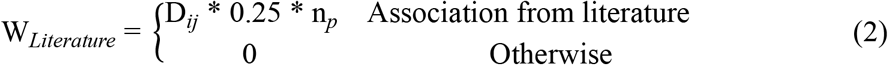

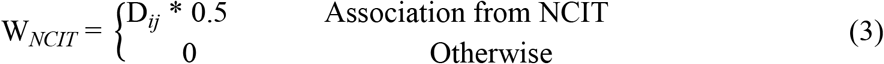

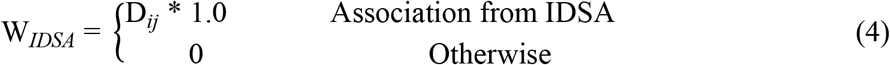

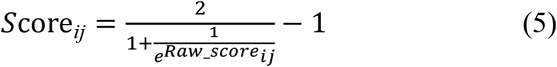

**Table 1.** The weights of different evidential sources according to their reliabilities

| Evidence | Evidence definition | Weight |
| --- | --- | --- |
| Literature | Author statement supported by traceable literature used in manual assertion | 0.25 |
| Data resource | Statement supported by manual curate data resource such as NCIT | 0.5 |
| Guideline | Consensus statement supported by the IDSA guideline | 1.0 |

Finally, the sigmoid function was used to normalize *Raw_score_ij_* to limit the range of the final association score *Score_ij_* from −1 to 1. In Equation (5), ‘*e*’ represents the natural constant *e*. *Score_ij_* can be used to judge the confidence of the relationship between a microbe and a disease phenotype. Please see the score distribution in **Figure 2**. A *Score_ij_* more than 0 indicates that the occurrence of the disease correlates with an increase of the microbial abundance, and a *Score_ij_* less than 0 indicates that the occurrence of the disease correlates with a decrease of the microbial abundance. The greater the absolute value of *Score_ij_*, the higher the number of previous reports of the respective microbe-disease association; the closer the score is to zero, the lower the number of previous reports of the respective microbe-disease association. By investigating the *Score_ij_* distribution, most associations were found with *Score_ij_* between −0.3 and 0.3 and the two peaks with *Score_ij_* more than 0.3 were involved in high confidence associations from NCIT and IDSA (Figure 2). This suggested that the score points of −0.3 and 0.3 would be the highly reliable thresholds to assess the confidence level of an association.

**Figure 2.**
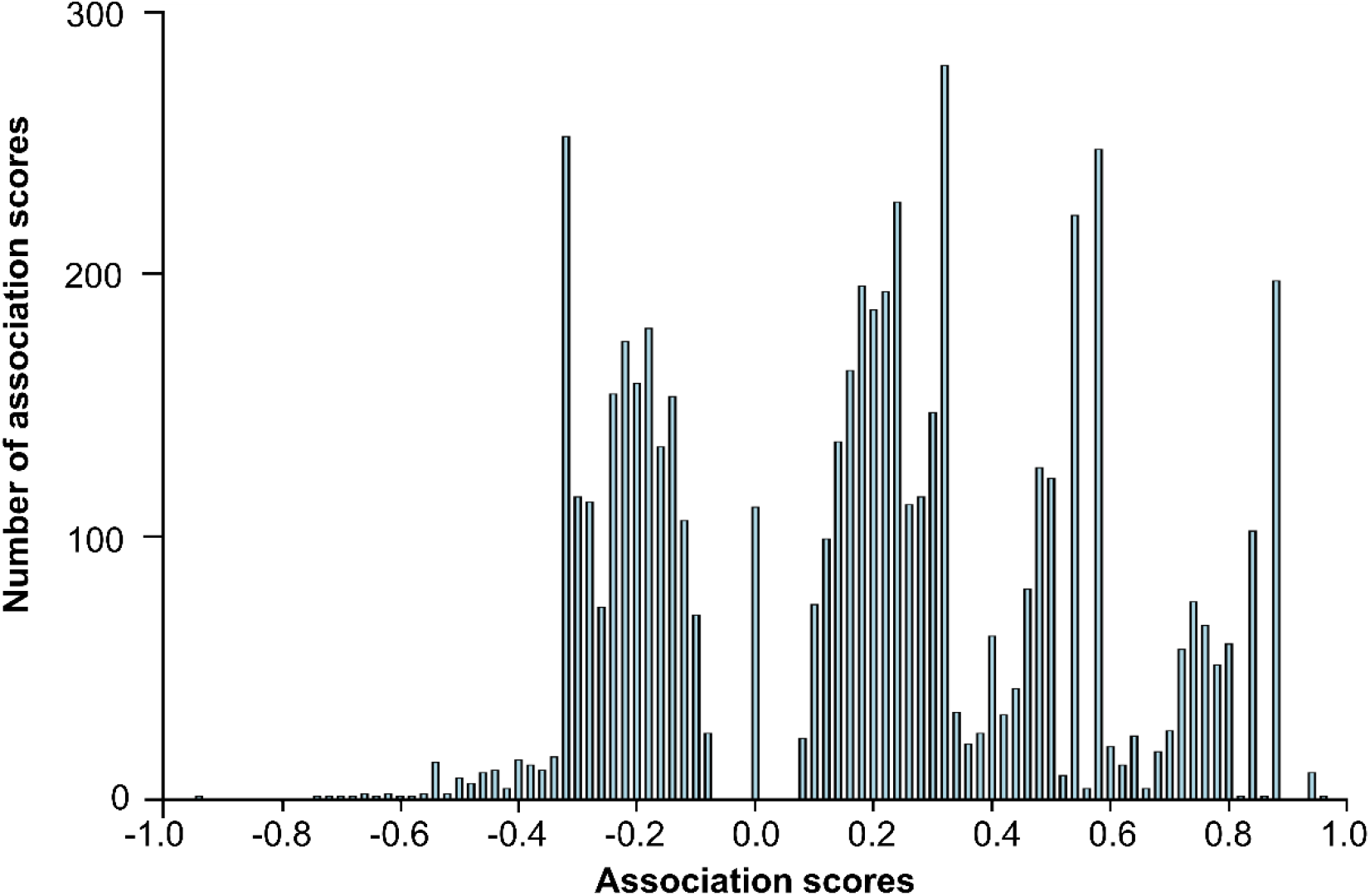
Data score distribution in MicroPhenoDB

### Implementation

The web applications in MicroPhenoDB were implemented in Java language by using the model-view-controller model and the SpringBoot framework, and were deployed on an Apache Tomcat web server. The association data of microbes and disease phenotypes were stored in a MySQL database. Data access, search, and visualization were implemented by using the Ajax API technology. The frontend interface was visualized by using the Vue.js framework. The sequence search tool was implemented using the EMBL-EBI tool framework [35].

## Database content and usage

### Database content

MicroPhenoDB collated 7449 redundant associations between 1781 microbes and 542 human disease phenotypes across more than 22 human body sites (**Table 2**). Of the 7449 associations, 29.7% were manually curated from the IDSA guideline (1196, 16.1%) [22], NCIT (849, 11.4%) [23], and peer-reviewed publications for human respiratory infection virus (164, 2.2%), and the others were consistently integrated with annotation from HMDAD (673, 9.0%) [18] and Disbiome (4567, 61.3%) [19] (**Figure 3A**). Multiple publications might support the same association between a microbe and a disease phenotype. After removing data redundancy based on the supporting publications, MicroPhenoDB produced 5677 non-redundant microbe-disease phenotype associations (Table 2). The number of non-redundant associations was over 11-fold (5677/483) of that in HMDAD. Each non-redundant association was assigned with a unique accession number (e.g. MBP00000900) and an association score. For the microbe distribution, MicroPhenoDB contained 1497 bacteria in a broad sense (including 1474 bacteria in a narrow sense, 11 *Rickettsia*, 6 *Chlamydia*, 4 *Ehrlichia*, and 2 *Mycoplasma*), 183 viruses, 58 fungi, and 43 parasites (Table 2). Approximately 88.3% (5014/5677), 8.5% (481/5677), 2.0% (116/5677), and 1.2% (66/5677) of the associations were related to bacteria, viruses, fungi, and parasite respectively (Figure 3B). The top six frequent disease-associated bacteria phyla were firmicutes, proteobacteria, bacteroidetes, actinobacteria, spirochaetes, and fusobacteria. The top disease-associated fungal phylum was ascomycota. Firmicutes included 271 genus/species in 4 classes (bacilli, clostridia, erysipelotrichia, and negativicutes) (Figure 3C). The microbes were mainly distributed in the body sites of gastrointestinal tract (37.3%), oral cavity (9.5%), respiratory tract (6.9%), skin and soft tissue (4.2%), urinary tract (3.5%), vagina (2.5%), and central nervous system (2.0%) (**Table 3**). The disease phenotypes were classified and described by EFO [30]. Many diseases were associated with pathogenic microorganisms, such as bacterial, digestive system, nervous system and autoimmune diseases (Figure 3D).

**Table 2.** Data scopes and scales in MicroPhenoDB

| Data scope | Data scale |
| --- | --- |
| Association | 5677 non-redundant microbe-disease associations. |
| Microbe | 1781 microbe species including 1497 bacteria in a broad sense (including 1474 bacteria in a narrow sense, 11 *rickettsia*, 6 *chlamydia*, 4 *ehrlichia*, and 2 *mycoplasma*), 183 viruses, 58 fungi and 43 parasites. |
| Disease phenotypes | 542 disease phenotypes annotated with EFO across more than 22 body sites. |
| Core genes | 685 bacteria and viruses annotated with 27,277 unique clade-specific core genes. |
| Virulence factors | 65 (4.3% of 1497) of bacteria annotated with more than 4204 virulence factor information, including pathogenic species, virulence factor gene name, characteristics structure, pathogenic mechanism, etc. |
| Antibiotic resistance data | 66 (4.4% of 1497) of bacteria annotated with more than 2522 antimicrobial resistance information, including resistance genes, resistance mechanisms, related antibiotics, etc. |

**Figure 3.**
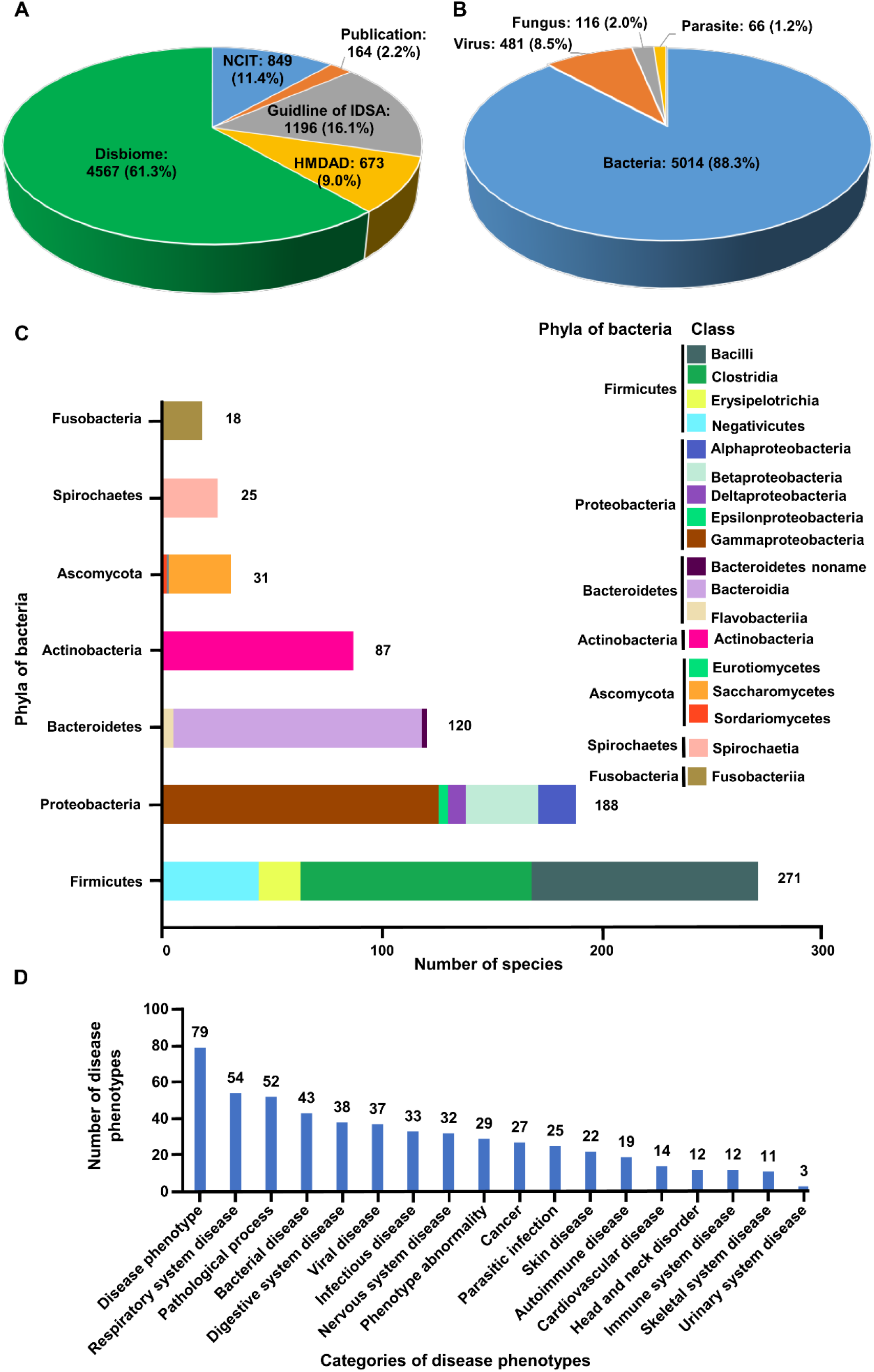
Data content and distribution in MicroPhenoDB. **A.** The association data collected from different resources. **B.** The distribution of different microbe types. **C.** The number of bacterial species in different phyla. **D.** The disease distribution in MicroPhenoDB.

**Table 3.** The top ten body sites of disease-associated microbes in MicroPhenoDB

| Body position | Association number(n) | Percent(n%) |
| --- | --- | --- |
| Gastrointestinal tract | 2119 | 37.3% |
| Oral cavity | 542 | 9.5% |
| Respiratory tract | 391 | 6.9% |
| Skin and soft tissue | 239 | 4.2% |
| Urinary tract | 197 | 3.5% |
| Vagina | 143 | 2.5% |
| Central nervous system | 114 | 2.0% |
| Nasal cavity | 85 | 1.5% |
| Bloodstream | 83 | 1.5% |
| Throat | 69 | 1.2% |

In total, 27,277 unique clade-specific core genes of 685 bacteria and viruses were retrieved from the dataset in MetaPhlAn2 and were annotated with gene functions using InterProScan (Table 2). In addition, 4204 virulence factor genes and 2522 drug resistance genes were also included from VFDB [20] and CARD [21], respectively. A small percentage ((4.3%, 65/1497) and (4.4%, 66/1497)) of bacteria was annotated with virulence factor information and antimicrobial resistance information, respectively (Table 2).

### Web interface

The MicroPhenoDB website (http://www.liwzlab.cn/microphenodb) provides user-friendly web interfaces to enable users to search, browse, prioritize, and analyze the microbe-disease association data in the database (**Figure 4**). The website offers multiple optional search applications of microbes, diseases, and associations to acquire prioritized association data with body site and microbe type filters. The prioritized microbe-disease associations can be downloaded as a CSV file for further analysis. The hierarchical structure of microbes and diseases are respectively displayed in the ‘Browse’ web page. Information regarding the increasing or decreasing tendency of microbial abundance in a disease, virulence factor, and antibiotic resistance of the microbes along with its core gene information are available in the ‘Browse’ web page. In addition, MicroPhenoDB provides the web service APIs for programmatical access of the association data and produces an output in the JSON format. All the association data and the API documentation are available on the website. Users are also encouraged to submit their data of newly published microbe-disease associations. Once checked by our professional curators and approved by the submission review committee, the submitted record will be included in an updated release.

**Figure 4.**
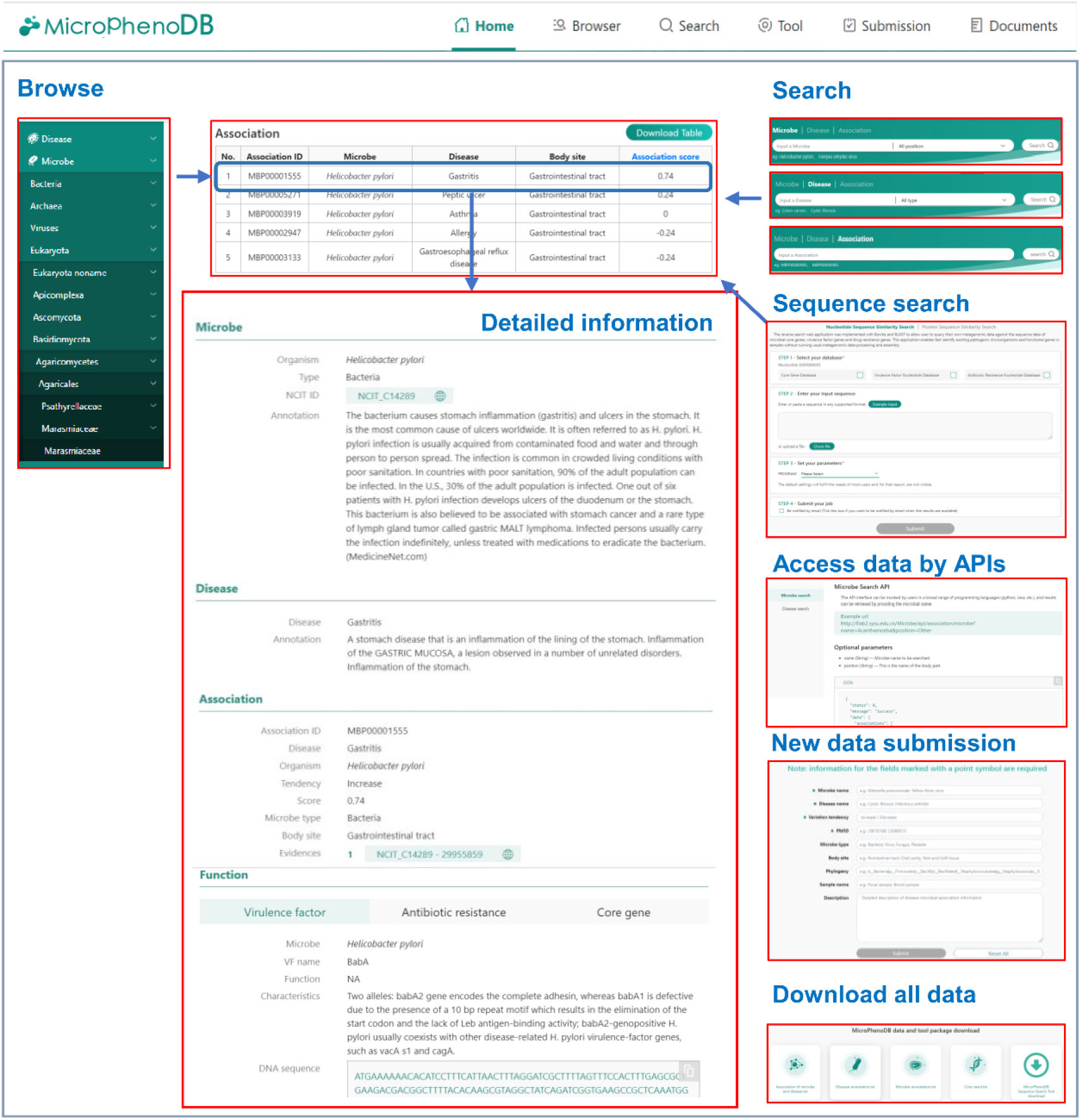
The MicroPhenoDB web interface

### Applications of association data

#### MicroPhenoDB sequence search to explore metagenomics data

In MicroPhenoDB, microbes were connected with diseases through 5677 non-redundant associations and linked to unique clade-specific core genes via 696,934 relationships (**Figure 5**). Core genes could serve as a hub to connect metagenomic sequencing data to microbes and their associated diseases (Figure 5). A sequence search application was implemented on the MicroPhenoDB website (http://www.liwzlab.cn/microphenodb/#/tool) to allow users to query their own metagenomic sequencing data against the MicroPhenoDB sequence datasets through the sequence alignment tools BLAST [36] and Bowtie2 [37] (Figure 5). The application can directly identify the composition of pathogenic microorganisms in metagenomic samples and can suggest potential disease phenotypes that may be caused, without running the usual metagenomic sequencing data processing and assembly, which are both time and resource consuming. Functional annotation for microbial core genes by the application includes gene ontology and pathway information. Searching against the sequence datasets of microbial pathogenic factors and drug resistance genes allows the identification of homologous genes and proteins related to virulence factors and antibiotic resistance (Figure 5).

**Figure 5.**
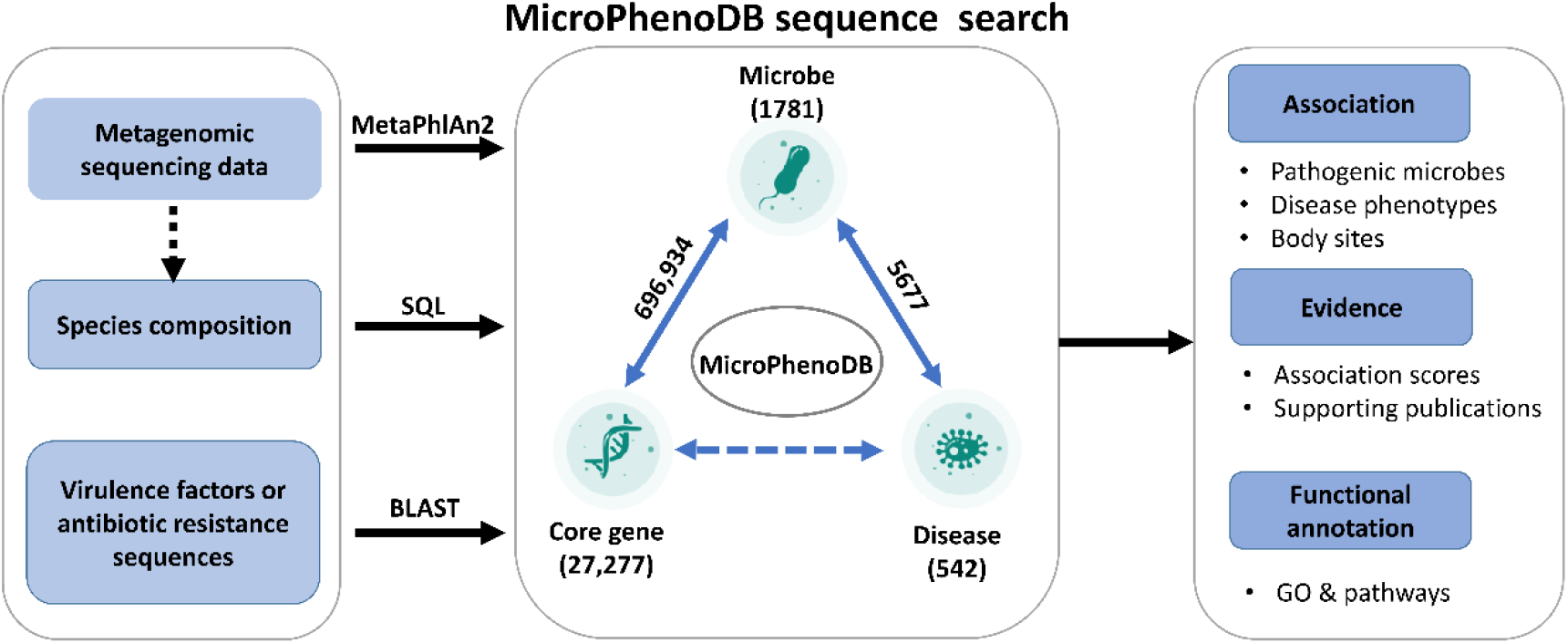
The MicroPhenoDB sequence search connects microbes, core genes and disease phenotypes

To assess the sequence search usability, we used the sequence search application to analyze an existing metagenomic dataset downloaded from the Genome Sequence Archive (accession: PRJCA000880) [38]. The dataset contained metagenomics data of lung biopsy tissues from 20 patients with pulmonary infection [39]. Our results identified pathogenic microbes in 95% (19 of 20) of patients, significantly higher than the 75% identification rate (15 of 20) found through the original metagenomic NGS (mNGS) analysis [39]. In addition, our search identified 37 pathogenic microbes in patients, while the mNGS method only identified 29 (Supplementary Table 2). Of the 37 microbes, 23 were identical to those by mNGS analysis. It was hard to estimate the false positives of the other 14 microbes, but we found that they may cause infections in patients with underlying diseases such as immunodeficiency. Therefore, this comparison suggested that the MicroPhenoDB sequence search application could screen metagenomic data for effective identification of pathogenic microbes. Due to the large size of metagenomic data and the need for a broadband network, we provide a software package of the search application for users to download and run locally. We also encourage users to upload the microbial abundance information to the online application for further analysis and visualization.

#### Identify unique clinical phenotypes of SARS-CoV-2 infection from different viral respiratory infections

The single-stranded RNA coronavirus SARS-CoV-2 can infect humans and cause COVID-19 disease [40]. Its structure is similar to those of viruses causing severe acute respiratory syndrome (SARS) and Middle East respiratory syndrome (MERS) [41]. At present, the diagnosis of SARS-CoV-2 infection is mainly based on clinical phenotypes, chest computed tomography (CT), and nucleic acid testing. Compared with CT and nucleic acid testing, clinical phenotype monitoring has significant advantages, such as a short turnaround time, low cost, and convenience [42]. To identify unique clinical phenotypes of SARS-CoV-2 infection from different viral respiratory infections, we searched MicroPhenoDB and obtained association data that contained 63 disease phenotypes and 14 respiratory tract infection viruses, such as human rhinovirus, parainfluenza virus, respiratory syncytial virus, metapneumovirus, and coronaviruses. The data were then imported into the Cytoscape software [43] for a network analysis. The output network (**Figure 6**) indicated that SARS-CoV-2 shares the clinical phenotype of pneumonia with the majority of other respiratory infection viruses, and the common clinical phenotypes of dry-cough, headache, fever, myalgia, vomiting, diarrhea and respiratory disease syndrome (underlined in green) with several influenza viruses and other coronaviruses. Importantly, the network also showed that dyspnea, fatigue, lymphopenia, anorexia, and septic shock (underlined in blue) were unique clinical phenotypes of SARS-CoV-2 infection compared with other viral respiratory infections [12, 44, 45]. Bear in mind that these phenotypes of SARS-CoV-2 infection might be frequent complications of other diseases and treatments. For example, dyspnea is a frequent complication of chronic respiratory diseases [46], lung cancer [47], and hepatopulmonary syndrome [48]; septic shock is a complication of pneumococcal pneumonia, chronic corticosteroid treatment, and current tobacco smoking [49]; fatigue is a complication of multi-type cancers [50, 51] and Parkinson’s disease [52]; and lymphopenia is a complication of human immunodeficiency viral infection [53]. However, our results suggest that these unique clinical phenotypes could help to identify SARS-CoV-2 infection from infections by SARS-CoV, MERS-CoV and other respiratory viruses.

**Figure 6.**
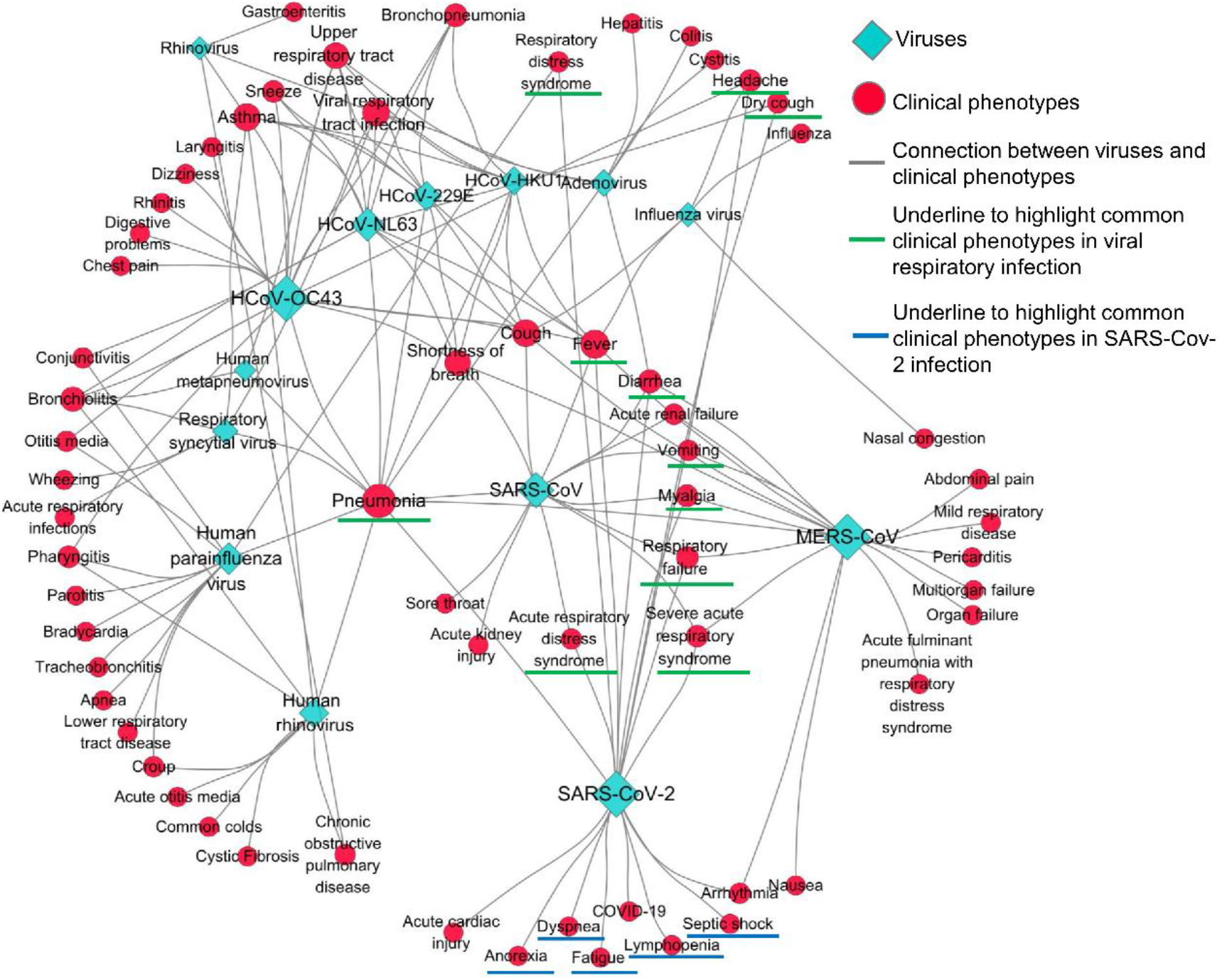
The Cytoscape network illustrates different clinical phenotypes across different viral respiratory infections

The diamonds represent the respiratory infection viruses. The red circles represent the disease phenotypes. Lager size of a circle or a diamond indicates more connections to a disease phenotype or a virus. The solid connection lines represent the associations between disease phenotypes and viruses with increasing abundance. Underlines indicate the clinical phenotypes discussed in the main text.

#### Association network in different body sites

The microbe-disease association data can be downloaded and used for further analysis. To generate a network to explore the reliable connections between the microbial changes and the diseases in multiple body sites, we obtained the association data of body sites such as vagina, urinary tract, and genitals using the reliable association score thresholds mentioned above (> 0.3 and < −0.3). The resulting association data were imported into the Cytoscape software [43] for network analysis. The output network (**Figure 7**) indicated that the decreasing abundance of *Lactobacillus* (underlined in red) was related to vaginal inflammation and bacterial vaginosis in vagina, while the increasing abundance of *Chlamydia* (underlined in green) resulted in lymphogranuloma venereum in genitals. Moreover, the network showed that the increasing abundance of *Mycoplasma genitalium* (underlined in blue) was associated with multiple diseases, which involve genitals, such as pelvic inflammatory disease, nongonococcal urethritis, and nonchlamydial nongonococcal urethritis. Furthermore, the network showed that a microbe abnormality could be associated with diseases involving different body sites. For example, the increasing abundance of *Neisseria gonorrhoeae* (underlined in purple) was associated with two diseases each in genitals and urinary tract. For users to assess the microbial pathogenicity, it is recommended to filter the data by using the association scores and follow the supporting publications for further investigation. User can follow our step-by-step guideline on the website (http://www.liwzlab.cn/microphenodb/#/guideline) to perform similar association analyses and generate Cytoscape networks.

**Figure 7.**
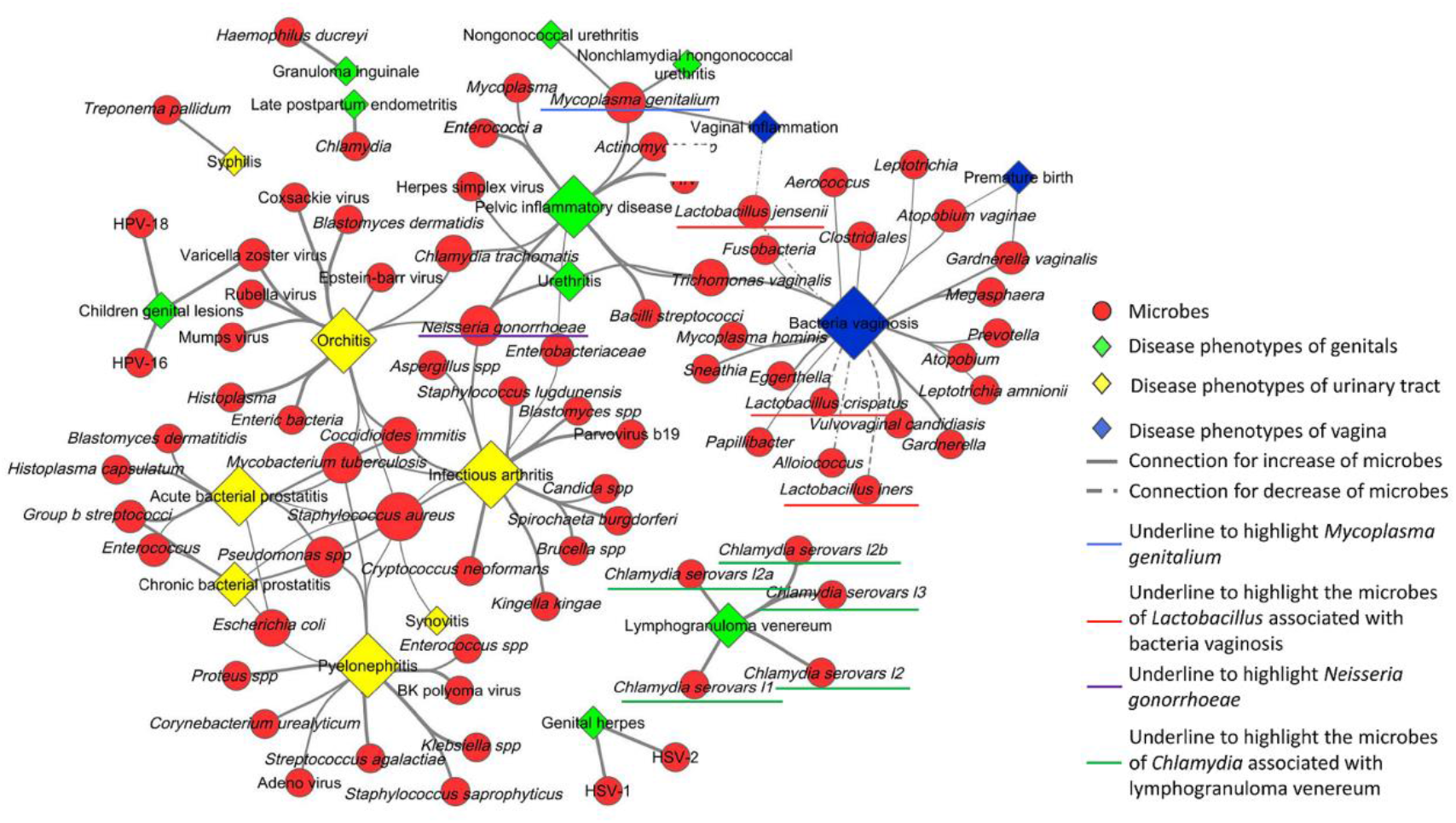
The Cytoscape network illustrates the associations between disease phenotypes and microbes at different body sites

The diamonds represent diseases resulted from a microbial abnormality at different body sites. The red circles represent the microbes. Lager size of a circle or a diamond indicates more connections to a disease phenotype or a virus. The solid connection lines represent the associations between diseases and microbes with increasing abundance, and the dash connection lines represent the associations between diseases and microbes with decreasing abundance. Underlines indicate the microbes discussed in the main text.

### Concluding remarks

Microbes play important roles in human health and disease. The curation and analysis of microbe-disease association data are essential for expediting translational research and application. In this study, we developed the MicroPhenoDB database by manually curating and consistently integrating microbe-disease association data. As far as we are aware, MicroPhenoDB is the first database platform to detail the relationships between pathogenic microbes, core genes, and disease phenotypes. In terms of data coverage, scoring models, and web applications, MicroPhenoDB outperformed data resources that contain similar association data (**Table 4**). For example, the numbers of associations, microbes, disease phenotypes, and supporting evidences in MicroPhenoDB were approximately 11.1, 6.1, 13.9, and 18.9-fold of those in HMDAD, respectively. Compared with both HMDAD and Disbiome, MicroPhenoDB refined the confidence scoring model using extra evidential metrics with different weights; it standardized the association annotations by manual curation, and included pathogenic data of virulence factors, microbial core genes, and antibiotic resistance gens. Moreover, MicroPhenoDB implemented the web applications and APIs for pathogenic microbe identifications in metagenomic data.

**Table 4.** Database contents and features of MicroPhenoDB compared with other databases

| **Data content & web applications** | **MicroPhenoDB** | **HMDAD** | **Disbiome** | **MicroPhenoDB/HMDAD** | **MicroPhenoDB/Disbiome** |
| --- | --- | --- | --- | --- | --- |
| Association data | 7449 | 673 | 4567 | 11.1 | 1.6 |
| Microbe organism | 1781 | 292 | 1292 | 6.1 | 1.4 |
| Microbe standardized annotation | 1041 | - | - | - | - |
| Organism taxonomy | 1032 | - | - | - | - |
| Disease phenotype | 542 | 39 | 282 | 13.9 | 1.9 |
| Disease standardized annotation | 446 | - | - | - | - |
| Supporting evidence | 1150 | 61 | 822 | 18.9 | 1.4 |
| Association score | Yes | None | None | - | - |
| Virulence factor | 4204 | - | - | - | - |
| Antibiotic resistance gene | 2522 | - | - | - | - |
| Core gene | 696,934 | - | - | - | - |
| Sequence alignment | Yes | None | None | - | - |
| Identification of pathogenic microbe | Yes | None | None | - | - |
| Web service API | Yes | None | None | - | - |

In MicroPhenoDB, many associations with confident scores came from our manual curation of the up-to-date clinical guidelines, which were supported by IDSA and ASM. MicroPhenoDB assigned higher weight values to the associations derived from the guidelines, and lower weight values to the associations from other literature data and databases. The original model for scoring confidence of the disease-microbe associations in HMDAD was based on a single literature evidence. Our MicroPhenoDB score model rated different supporting evidence according to the credibility of related sources and provides a score to evaluate a disease-microbe association.

By integrating unique, clade-specific microbial core genes and using the data from MetaPhlAn2, the MicroPhenoDB sequence search application enables rapid identification of existing pathogenic microorganisms in metagenomic samples without running the usual sequencing data processing and assembly. However, the resulting associations from the sequence search do not guarantee microbial pathogenicity, but provide clues for further investigation. The annotated core genes are also limited in size and cannot represent all microbial species. To consistently analyze the important functions of microbes, other data or tools are also recommended, such as UniRef clusters [54], MetaCyc [55], HUMAnN2 [56], and pan-genomic data.

To serve the research community, we will update the database every six months and constantly improve it with more features and functionalities. As a novel and unique resource, MicroPhenoDB connects pathogenic microbes, microbial core genes, and disease phenotypes; therefore, it can be used in metagenomic data analyses and assist studies in decoding microbes associated with human diseases.

### Data availability

To access the association data, the online applications and the software package, please visit http://www.liwzlab.cn/microphenodb/#/download.

## Authors’ contributions

GY and WZ performed the majority of the study including data curation, scoring model implementation, web development and manuscript preparation. MY managed the web server. HY and JW contributed to the scoring model design. HZ, LW, ZX revised the manuscript and provided suggestions. WL designed and managed the overall study, as well as editing the manuscript.

## Acknowledgements

This work was supported by the National Key R&D Program of China [grant number 2016YFC0901604], the National Natural Science Foundation of China [grant number 31771478] and the National Key R&D Program of China [grant number 2018YFC0910401] to WL.

## Supplementary material

**Supplementary Figure 1.**
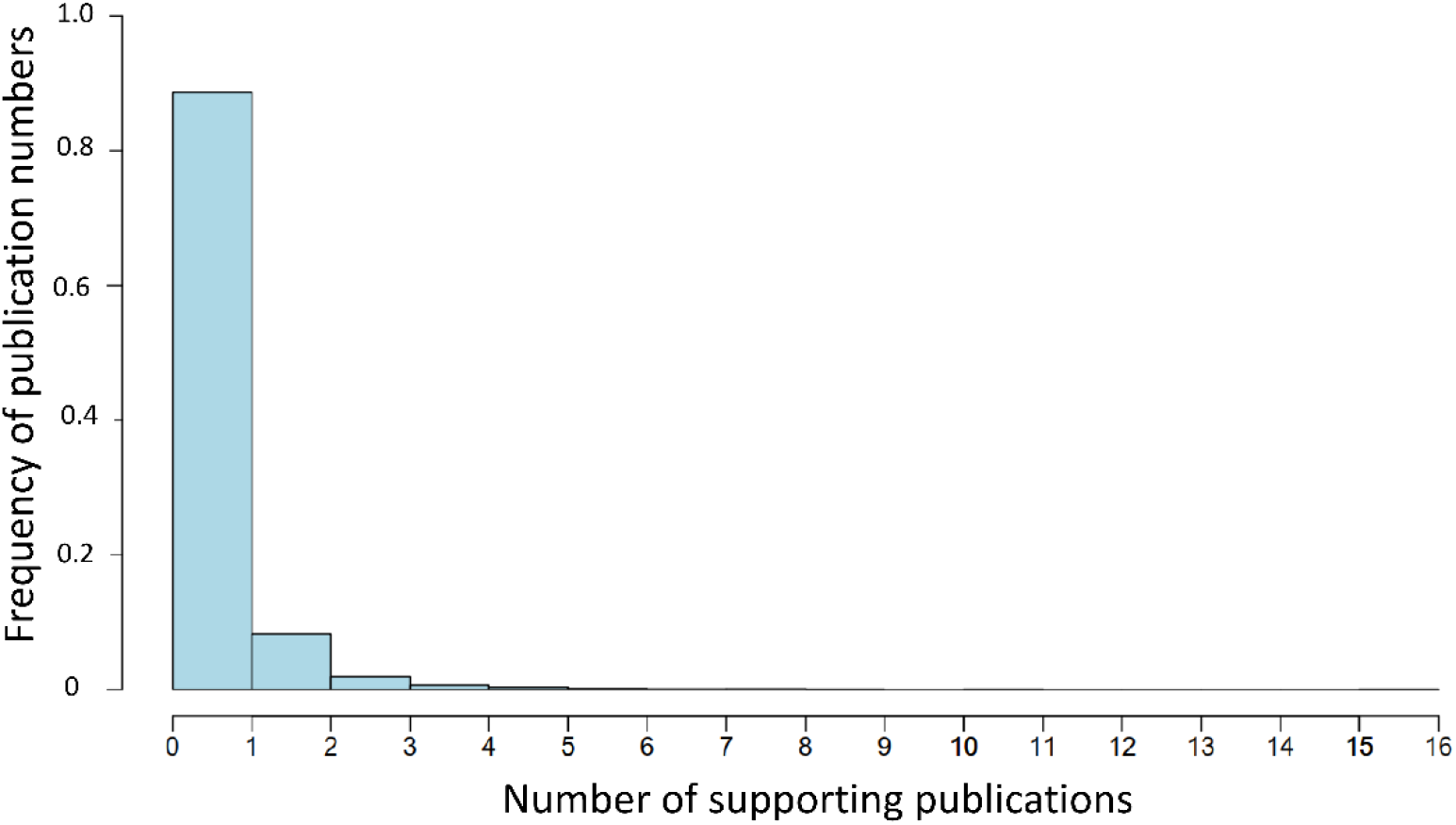
The distribution of numbers of supporting publications. The blue histogram represents the frequency of the number of supporting publications.

**Supplementary Table 1.** The version or release of databases and tools used in the MicroPhenoDB construction

| Name | URL | Date | Version | Ref |
| --- | --- | --- | --- | --- |
| HMDAD | http://www.cuilab.cn/hmdad | 2016 | Version 1.0 | [18] |
| Disbiome | https://disbiome.ugent.be/ | 2018 | Version 1.0 | [19] |
| IDSA guideline | https://pubmed.ncbi.nlm.nih.gov/30169655/ | 2018 | 2018 update | [22] |
| EFO | https://www.ebi.ac.uk/efo/ | 2019 | Version 3.18.0 | [30] |
| NCIT | https://www.ebi.ac.uk/ols/ontologies/ncit | 2019 | Version 19.11d | [23] |
| VFDB | http://www.mgc.ac.cn/VFs/ | 2016 | Version 1.0 | [20] |
| CARD | http://arpcard.mcmaster.ca | 2017 | Version 1.0 | [21] |
| MetaPhlAn2 | http://huttenhower.sph.harvard.edu/metaphlan2/ | 2015 | Version 2.0 | [28] |
| InterProScan | http://www.ebi.ac.uk/interpro/ | 2019 | Version 72.0 | [29] |
| NCBI-taxonomy | https://www.ncbi.nlm.nih.gov/taxonomy | 2019 | - | [26] |
| EBI tool framework | https://www.ebi.ac.uk/ | 2019 | - | [35] |
| Genome Sequence Archive | http://gsa.big.ac.cn | 2020 | - | [38] |

**Supplementary Table 2.**
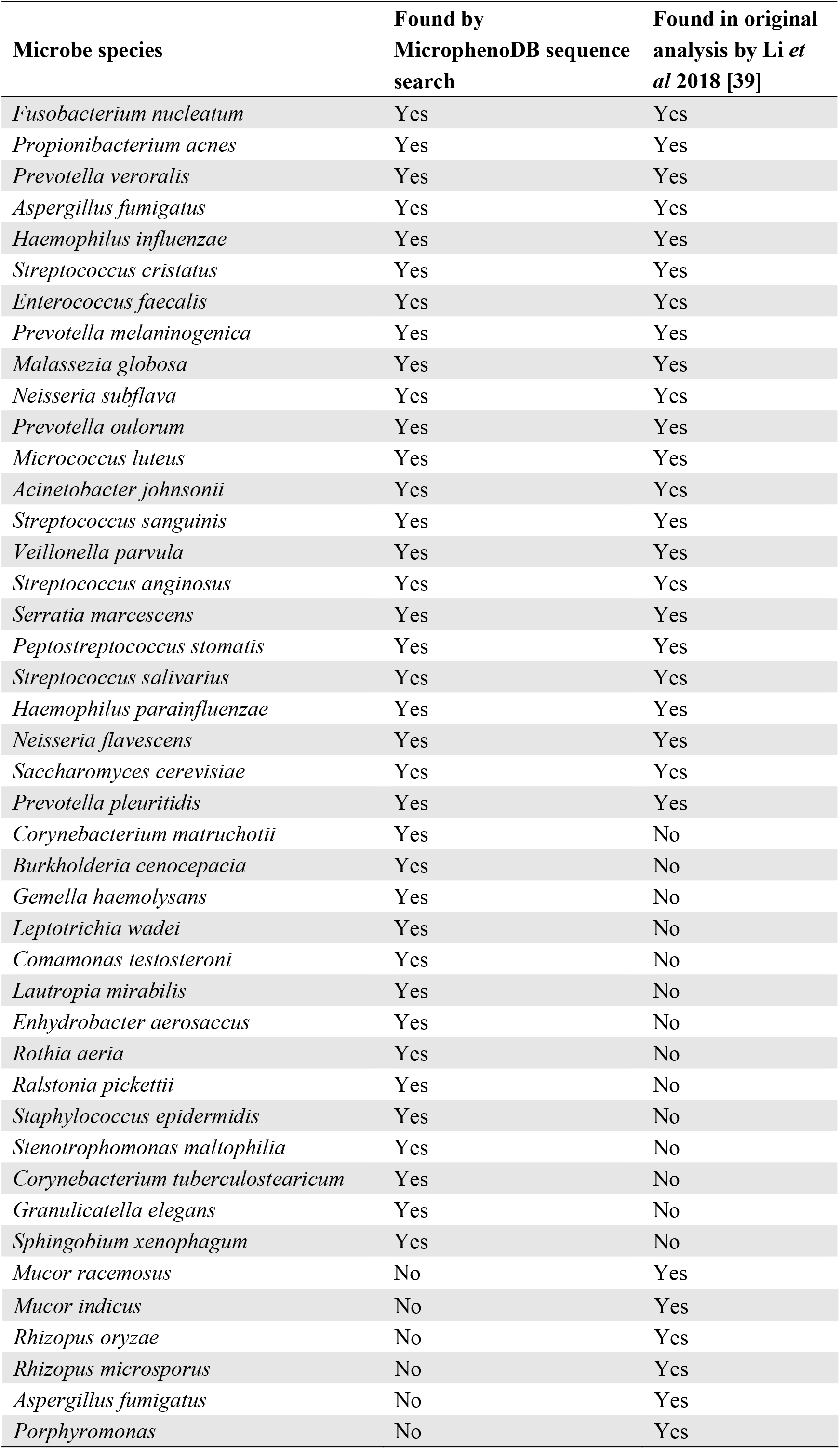
The analysis result by MicroPhenoDB sequence search in an existing metagenomic dataset

